# Hierarchies & Lower Bounds in Theoretical Connectomics

**DOI:** 10.1101/559260

**Authors:** Venkatakrishnan Ramaswamy

## Abstract

Connectomics is a sub-field of Neuroscience aimed at determining connectomes – exact structures of neurons and their synaptic connections in nervous systems. A number of ongoing initiatives at the present time are working towards the goal of ascertaining the connectomes or parts thereof of various organisms. Determining the detailed physiological response properties of all the neurons in these connectomes is out of reach of current experimental technology. It is therefore unclear, to what extent knowledge of the connectome alone will advance a mechanistic understanding of computation occurring in these neuronal circuits, especially when the high-level function(s) of the said circuit is unknown.

We are pursuing a research program to build theory in order to investigate these issues. In previously published work [1], towards this end, we have developed a theory of connectomic constraints for feedforward networks of neurons. Specifically, for feedforward networks equipped with neurons that obey a deterministic spiking neuron model, we asked if just by knowing the structure of a network, we could rule out spike-timed computations that it could be doing, no matter what response properties each of its neurons may have. Our neurons were abstract mathematical objects that satisfied a small number of axioms that correspond to certain broadly-obeyed properties of neurons.

Here, we develop additional theoretical tools and notions to address these questions. The idea is to study the space of all possible spike-train to spike-train transformations. We are interested in asking how the subset of transformations spanned by networks of specific architectures can be related to hierarchical subsets of the space that are characterized by particular mathematical properties of transformations. In particular, given such a hierarchy of subsets, what is the “smallest” subset that contains the set of transformations spanned by networks of a specific class of architectures? Even if one cannot establish such a subset exactly, proving bounds on it (according to the hierarchy) might offer insight. After setting up the mathematical framework to make these notions precise, we construct explicit classes of hierarchies and prove a number of such lower bounds.

## 1 Introduction

Progress in experimental techniques, since the turn of the century [2, 3, 4, 5, 6, 7], has led to a rekindling of interest in determining exact structures of (microscale) neural circuits that comprise nervous systems [8, 9, 10, 11, 12], which were rechristened as connectomes [13]. The first connectome – that of the nematode *Caenorhabditis elegans* – was determined in 1986 [14], after an effort that took over a decade, much of it manual. The only other complete connectome to be determined, to date, is that of the larval tadpole *Ciona intestinalis* [15], which has an asymmetric nervous system and in fact has fewer neurons than *C. elegans*; this was accomplished using modern automated techniques. That said, a significant percentage of the larval Drosophila connectome has already been reconstructed, although it is not publicly available at the time of writing [16]. Likewise, the electron microscopy volume of the entire larval zebrafish [17] and adult Drosophila [18] are publicly available, although they haven’t been fully reconstructed at the time of writing. Indeed, a number of studies have already fruitfully used Connectomics to address several questions about neural circuits and their function [19, 20, 12, 21, 22, 23, 24, 25, 26, 27, 28]. Also, it has been possible to first perform functional imaging of neural circuit activity in-vivo and then perform connectomic reconstruction of circuits in the same piece of tissue [19, 20, 28].

Current techniques, however, do not allow determination of detailed physiological response properties of all neurons in the connectomes and this is likely to be the case in the foreseeable future. It is therefore unclear to what extent knowledge of the connectome will inform us about mechanistic computation occuring in the neural circuits being reconstructed. Indeed, this has been one criticism of the connectomics efforts [29], in that it is unclear what exactly we will learn by knowing the connectome. Part of the criticism originates from a lack of theory. In particular, there is a need for theory that can use connectomic data to infer constraints on computation in neural circuits and generate hypotheses about mechanistic computation in the neural circuits. We are pursuing a research program to build such theory. To this end, in previously published work [1], we have developed a theory of connectomic constraints on computation in neural circuits. Specifically, for feedforward networks equipped with neurons that obey a deterministic spiking neuron model, we asked if just by knowing the structure of a network, we can rule out computations that it could be doing, no matter what response properties each of its neurons may have. We also stipulated the need to demonstrate a network with a different structure comprising “simple” neurons that could indeed effect the computation in question. After setting up a mathematical framework within which these questions could be precisely posed, we showed results of this form (which we call *complexity results*) for certain classes of network architectures. We also proved, mathematically, that for certain other classes of network architectures, given our limited assumptions on the individual neurons^[1]^, there are fundamental limits to constraints imposed by network structure alone. Our neurons were abstract mathematical objects that satisfied a small number of axioms that correspond to certain broadly obeyed properties of neurons. Complexity results, thus, were in the form of mathematical proofs that use these axioms and the structure of the network in question to establish explicit spiketrain to spike-train transformations that could not be effected by any network of the said structure, which could in turn be effected by a network of a different structure. Indeed, the broad program in this line of research is to start from first principles, as we have done in [1] and prove such results for networks of axiomatic neurons with progressively larger number of axioms and also generalize the theory to treat the case of recurrent networks. The idea is to eventually use this type of theory as a starting point to rule out specific computations in neural circuits for which connectomic data is available. This might then be used to formulate hypotheses about mechanistic computation in such circuits, which could be tested experimentally. In addition to this goal, the theory would also allow us to understand which aspects of network structure are crucial to the manifestation of what kinds of spike-timed computation. This could, in principle, aid in understanding why some types of network motifs might be conserved across individuals or species, especially in the emerging field of Comparative Connectomics.

Here, we develop additional theoretical tools and notions to address these questions. The idea is to study the space of all possible spike-train to spike-train transformations. In particular, we are interested in asking how the subset of transformations spanned by networks of specific architectures can be related to subsets of the space that are characterized by particular mathematical properties of transformations. While the former types of subsets are related to networks of neurons, the latter type are related to transformations alone and mathematical properties thereof. More concretely, using mathematical properties of transformations (i.e. without any reference to neurons or networks), the idea is to identify a sequence of subsets of this space, with each subset contained in the subsequent one in the sequence. We call this sequence of subsets a *Transformation Hierarchy*. This allows us to relate sets in a transformation hierarchy to sets of transformations spanned by specific network architectures, which we call the *Complexity Classes* of those architectures. We do this by finding the “smallest” set in the hierarchy that contains the complexity class in question as a subset. This set is called the *Hierarchy Class* of the architecture with respect to the said transformation hierarchy. Figure 1 provides a Venn Diagram illustrating these notions. Even if we cannot establish the hierarchy class of a given architecture with respect to a hierarchy, proving bounds^[2]^ on them might offer insight. As an application of these notions, we have constructed explicit classes of transformation hierarchies. For every set (starting from a certain set) in each such hierarchy, we demonstrate network architectures, for which the set in question is a lower bound on the hierarchy class of the said network architecture.

**Figure 1.**
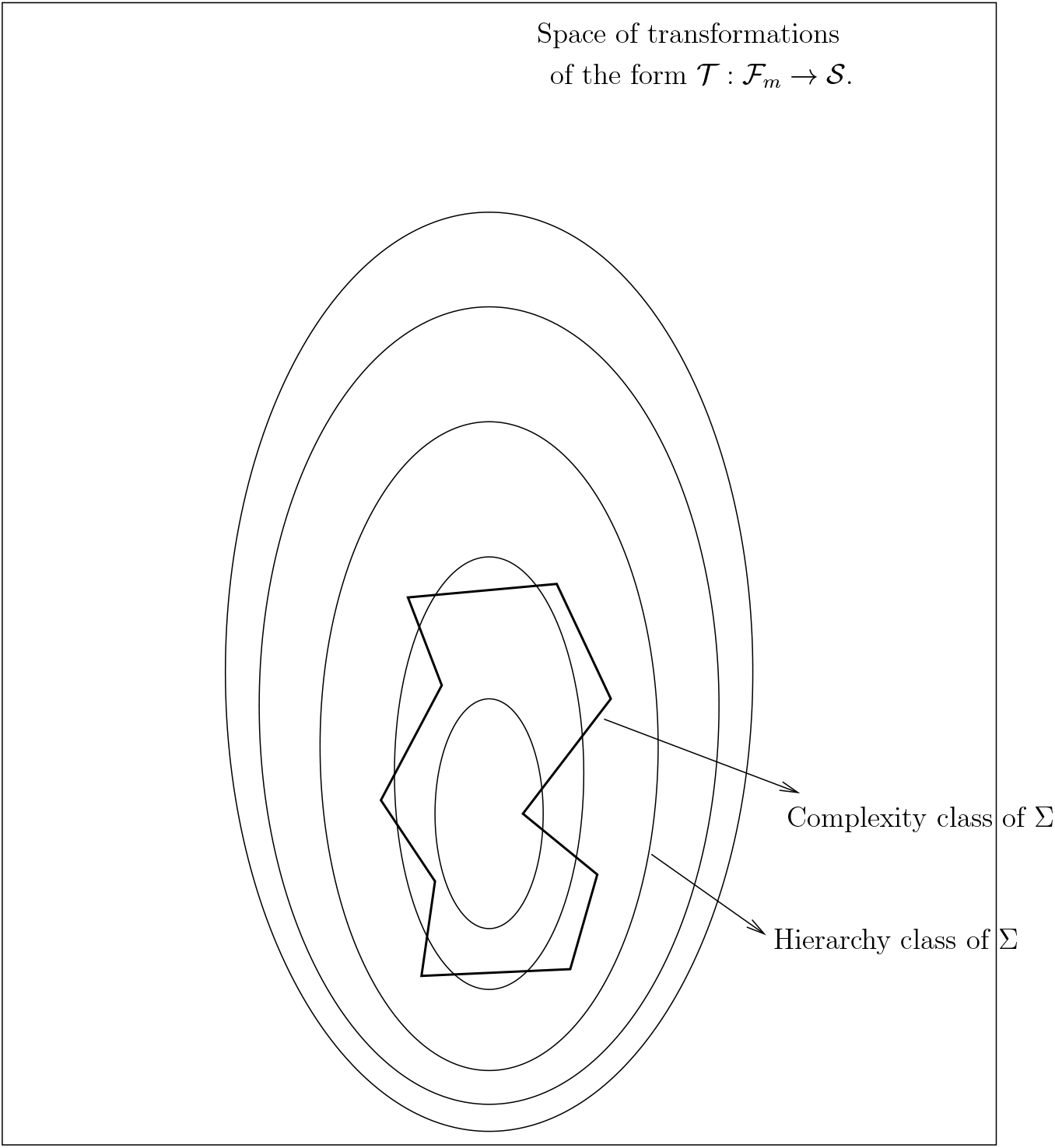
A schematic Venn Diagram illustrating the basic notions of a transformation hierarchy, complexity class and hierarchy class. Informally, a transformation hierarchy is a sequence of nested subsets in the space of transformations. The complexity class of a set of networks is the subset of that space spanned by transformation induced by networks in the said set. The hierarchy class of a set of network is the “smallest” set in the hierarchy that contains the complexity class as a subset.

The reader familiar with Theoretical Computer Science will observe that the approach taken here is somewhat reminiscent of that in Computational Complexity Theory, although the settings, details and questions are very different.

## 2 Definitions and Preliminaries

The treatment here is largely self-contained. In order to make this so, in this section, we reproduce verbatim from [1], definitions that constitute the basic mathematical formalism used to describe spike trains and operations on them.

An *action potential* or *spike* is a stereotypical event characterized by the time instant at which it is initiated in the neuron, which is referred to as its *spike time*. Spike times are represented relative to the present by real numbers, with positive values denoting past spike times and negative values denoting future spike times. A *spike-*train 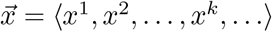 is a strictly increasing sequence of spike times, with every pair of spike times being at least *α* apart, where *α >* 0 is the absolute refractory period^[3]^ and *x*^*i*^ is the spike time of spike *i*. An *empty spike-train*, denoted by 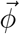, is one which has no spikes. A *time-bounded spike-train* (with *bound* (*a, b*)) is one where all spike times lie in the bounded interval (*a, b*), for some *a, b* ∈ ℝ. We use 𝒮 to denote the set of all spike trains and 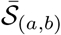 to denote the set of all time-bounded spike-trains with bound (*a, b*). A spike-train is said to have a *gap* in the interval (*c, d*), if it has no spikes in that time interval. Furthermore, this gap is said to be of *length d* − *c*.

We use the term *spike-train ensemble* to denote a collection of spike-trains. Thus, formally, a *spike-train ensemble* 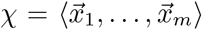 is a tuple of spike-trains. The *order* of a spike-train ensemble is the number of spike-trains in it. For example, 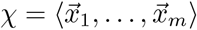 is a spike-train ensemble of order *m*. A *time-bounded spike-train ensemble* (with *bound* (*a, b*)) is one in which each of its spike-trains is time-bounded (with *bound* (*a, b*)). A spike-train ensemble *χ* is said have a *gap* in the interval (*c, d*), if each of its spike trains has a gap in the interval (*c, d*).

Next, we define some operators to time-shift, segment and assemble/disassemble spiketrains from spike-train ensembles. Let 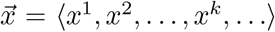 be a spike-train and 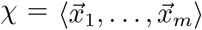 be a spike-train ensemble. The *time-shift operator for spike-trains* is used to time-shift all the spikes in a spike-train. Thus, 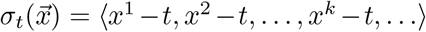. The *time-shift operator for spike-train ensembles* is defined as 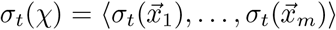.

The *truncation operator for spike-trains* is used to “cut out” specific segments of a spiketrain. It is defined as follows: 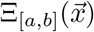 is the time-bounded spike-train with bound [*a, b*] that is identical to 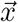 in the interval 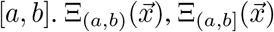 and 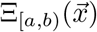) are defined likewise. In the same vein, 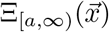 is the spike-train that is identical to 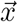 in the interval [*a,* ∞) and has no spikes in the interval (-∞, *a*). Similarly, 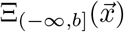 is the spike-train that is identical to 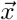 in the interval (-∞, *b*] and has no spikes in the interval (*b,* ∞). 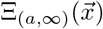 and 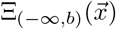 are also defined similarly. The *truncation operator for spike-train ensembles* is defined as 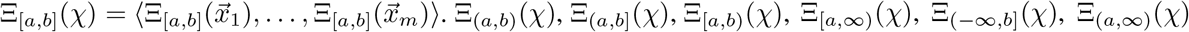 and Ξ _(-∞,*b*)_(*χ*) are defined likewise. Furthermore, Ξ _*t*_(·)is shorthand for Ξ _[*t,t*]_(·). The *projection operator for spike-train ensembles* is used to “pull-out” a specific spike-train from a spike-train ensemble. It is defined as 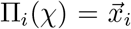, where 1 ≤ *i* ≤ *m*. Let 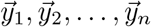 be spike-trains. The *join operator for spike-trains* is used to “bundle-up” a set of spike-trains to obtain a spike-train ensemble. It is defined as 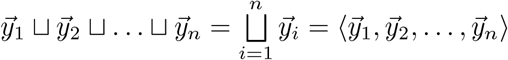

The neuron model used is a deterministic spiking neuron model. One example would be the abstract model used in [1], although a more restrictive model with more axioms would also allow for the notions defined in this paper.

Next, we have the notion of a Flush Criterion that appears as Definition 5 in [1].

### Definition (Flush Criterion)

A spike-train ensemble *χ* is said to satisfy a *T-Flush Criterion*, if all its spikes lie in the interval (0, *T)*, i.e. it has no spikes upto time instant *T* and since time instant 0.

Let the set of spike-train ensembles of order *m* that satisfy the T-Flush criterion be 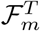 Let 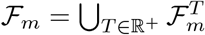.

Each feedforward network induces^[4]^ a transformation of the form 𝒯: ℱ_*m*_ → 𝒮, where 𝒮 is the space of all spike trains.

Next, we define the relation *more complex than*. For brevity, the definition below combines Definition 6 and Lemma 5 in [1] in order to define this relation equivalently in terms of spike-train ensembles satisfying the Flush Criterion.

### Definition (Transformational Complexity)

Let Σ_1_ and Σ_2_ be two sets of feedforward networks, each network being of order *m*, with Σ_1_ ⊆ Σ_2_. The set Σ_2_ is said to be *more complex than* Σ_1_, if there exists an 𝒩 ′ ∈ Σ_2_ such that for all 𝒩 ∈ Σ_1_, 𝒯_𝒩 ′_ ≠ 𝒯_𝒩_.

Each result of the above form is called a *complexity result*.

It is important to emphasize that we have skipped a great deal of other notation, notions, lemmas and theorems in [1] that are not central to the treatment here.

## 3 Complexity Classes, Transformation Hierarchies and Hierarchy Classes

Let 𝔉_*m*_ be the space of all possible transformations of the form 𝒯: ℱ_*m*_ → *S* that map spike spike-train ensembles of order *m* which satisfy the Flush criterion to output spike trains.Each acyclic network of order *m* induces one such transformation. A set of networks of order *m* therefore induces a class of such transformations, which we call the *complexity class* of that set.

### Definition 1 (Complexity Class)

Let Σ be a set of acyclic networks of order *m*. The *complexity class* of Σ, C_Σ_, is defined to be the set ⋃ 𝒩_∈Σ_𝒯_𝒩_.

As is clear from Lemma 6 in [1], no complexity class spans the entire space 𝔉_*m*_. Lemma 5 from [1] implies that questions of relative complexity of sets of networks can be posed in terms of questions about containment of their complexity classes. Figure 2 illustrates the situation and the next proposition formalizes it.

**Figure 2.**
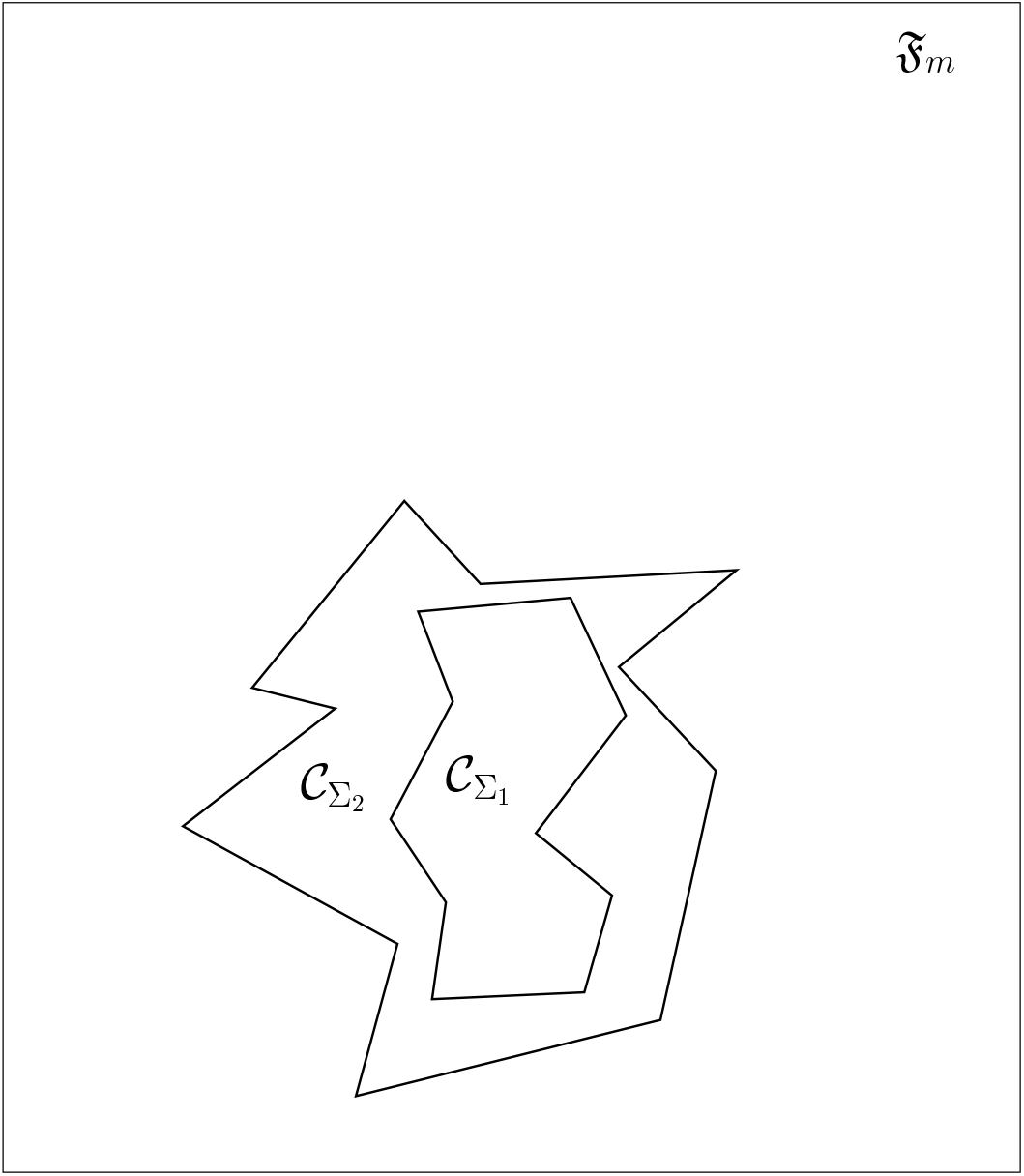
A schematic Venn Diagram showing complexity classes when one set of networks is more complex than another. If Σ_2_ is more complex than Σ_1_, it is equivalent to say that 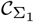 is a proper subset of 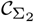.

### Lemma 1

*Let* Σ_1_ *and* Σ_2_ *be two sets of acyclic networks, each network being of order m, with* Σ_1_ ⊆ Σ_2_. *Further, let* 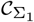 *and* 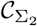 *be the corresponding complexity classes. Then,* Σ_2_ *is more complex than* Σ_1_ *if and only if* 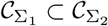

### Proof

First, suppose Σ_2_ is more complex than Σ_1_. That 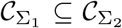 follows immediately from the fact that Σ_1_ ⊆ Σ_2_. Lemma 5 from [1] implies that ∃ 𝒩 ^′^ ∈ Σ_2_ such that ∀𝒩 ∈ Σ_1_ 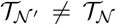 That is, 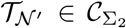 and 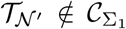 Since 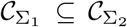, it follows that 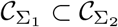.

To prove the other direction, assume 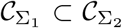 Therefore, ∃ 𝒯: ℱ_*m*_ → *𝒮* such that 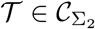 and 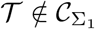. By definition of 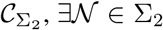, so that 𝒯_𝒩_ = 𝒯. Since 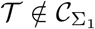, the definition of 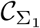 implies that ∀ 𝒩 ^′^ ∈ Σ_1_, 𝒯_𝒩′_≠ 𝒯. Therefore, Σ_2_ is more complex than Σ_1_.□

Next, we make precise the notion of a Transformation Hierarchy. Informally, a Transformation Hierarchy is a sequence of subsets of this space, with each subset contained in the subsequent one in the sequence.

### Definition 2 (Transformation Hierarchy)

A *Transformation Hierarchy* ℋ in 𝔉_*m*_ is a sequence of subsets ⟨*H*_1_, *H*_2_,*…, H_i_,…*, F_𝔉_⟩ of 𝔉_*m*_ with *H*_*i*_ ⊂ *H*_*i*+1_, ∀*i* = 1, 2,*…*.

Note that the above definition exists independent of the existence of networks. That is, a hierarchy is defined only in terms of properties of transformations in its constituent sets. Figure 3 provides an illustration. The next definition provides a connection between sets of acyclic networks and hierarchies. Each set of acyclic networks Σ is associated with a specific set in the hierarchy called its *hierarchy class*, so that the hierarchy class is the smallest set in the hierarchy that contains the complexity class of Σ.

**Figure 3.**
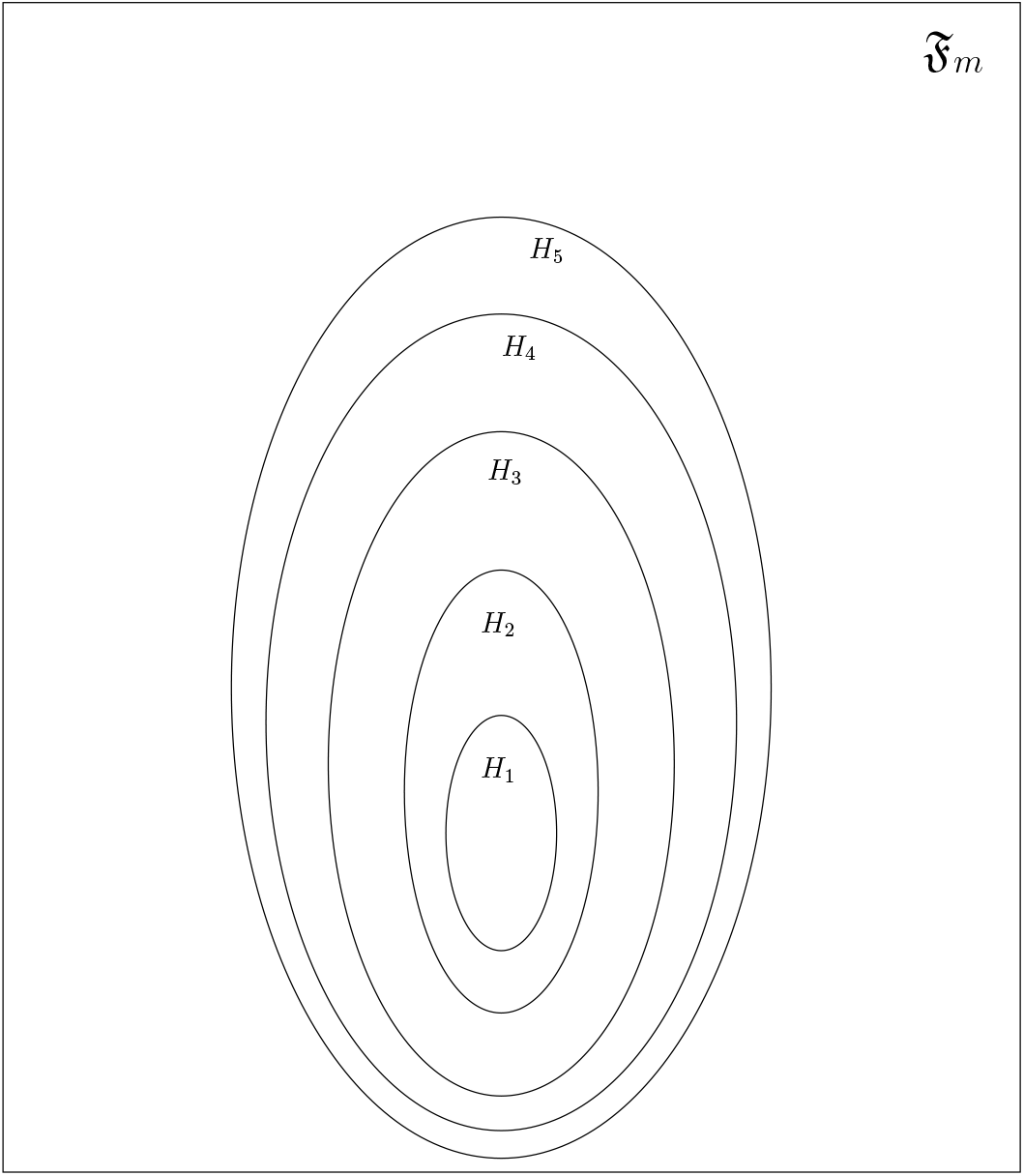
A schematic Venn Diagram illustrating a transformation hierarchy, which is a nested sequence of subsets of the space of transformations.

### Definition 3 (Hierarchy Class)

Let Σ be a set of acyclic networks, each of order *m*, and let ℋ = ⟨*H*_1_, *H*_2_,*…, H_i_,…,* 𝔉_m_⟩ be a transformation hierarchy in 𝔉_*m*_. The *Hierarchy Class H*_Σ_ of Σ in H is the set *H*_*i*_ with 𝒞_Σ_ ⊆ *H*_*i*_ and 𝒞_Σ_ ⊆ *H*_*i*-1_, if such an *H*_*i*_ exists and 𝔉_*m*_ otherwise.

Note that the Hierarchy class of a set of networks is well-defined. This is because 𝒞_Σ_ ⊂ 𝔉_*m*_, so there is atleast one set in the hierarchy that contains every complexity class. Also, the sets in the hierarchy are well-ordered^[5]^due to which every collection of sets from the hierarchy (which contain 𝒞_Σ_) has a smallest set.

The above definitions allow us to create a variety of hierarchies based on specific properties of transformations. If we can then say something about the hierarchy classes of specific architectures in each hierarchy, it enables us to get a better understanding of various aspects of the transformations effected by networks with this architecture.

Even if we cannot identify the hierarchy class of a given architecture in a hierarchy, proving bounds^[6]^ on them might give us some insight. Also, as alluded to before, these bounds can be used to establish that one set of networks is more complex than another.

A set in a hierarchy is an *upper bound* on a hierarchy class if it contains the hierarchy class as a subset. Likewise, a set in a hierarchy is a *lower bound* on a hierarchy class if the hierarchy class contains the set as a subset.

Bounds on hierarchy classes of specific sets of networks can be used to establish complexity results. If there are two sets of networks, the first contained in the second and if an upper bound on the hierarchy class of the first set is “smaller” than a lower bound on the hierarchy class of the second (with respect to the same hierarchy), then the second set is more complex than the first. Figure 4 gives a picture. Note that this is just a sufficient condition, not a necessary one, for one set to be more complex than the other^[7]^. The next lemma formalizes the above observations.

**Figure 4.**
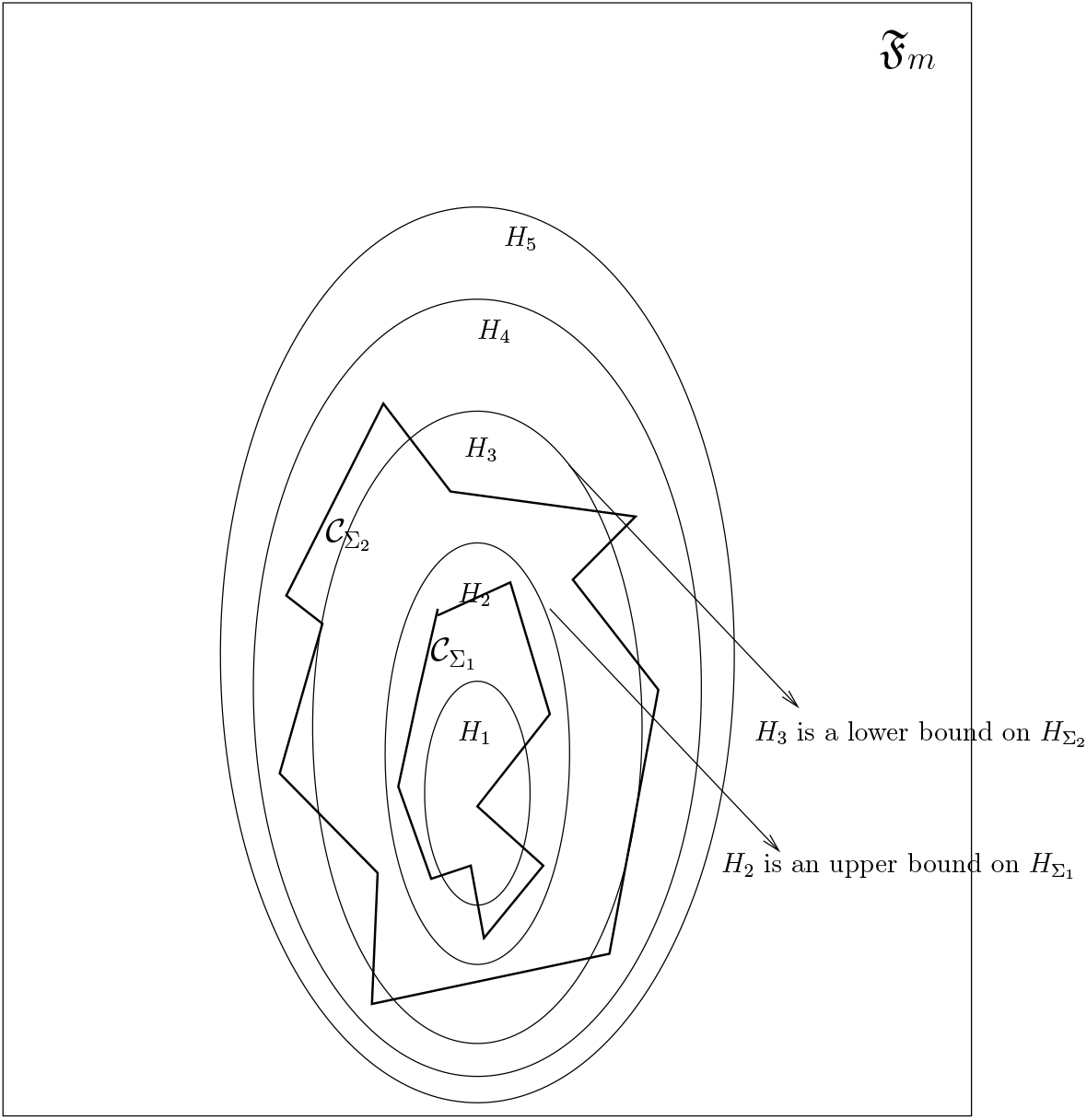
A schematic Venn Diagram demonstrating how upper bounds and lower bounds on hierarchy classes in a transformation hierarchy can be used to establish complexity results. In this case, the fact that an upper bound on 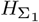 is a proper subset of a lower bound on 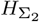 immediately implies that Σ_2_ is more complex than Σ_1_, as is proved in Lemma 2.

### Lemma 2

*Let* Σ_1_ *and* Σ_2_ *be two sets of acyclic networks, each comprising networks of order m, with* Σ_1_ ⊆ Σ_2_. *Furthermore, let* 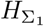 *and* 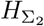 *be the corresponding hierarchy classes in a transformation hierarchy* ℋ = 〈*H*_1_, *H*_2_,…, *H*_*i*_,…, ℑ_m_〉 *in* ℑ_m_. *Moreover, let H_u_ be an upper bound on* 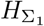 *and H_l_ be a lower bound on* 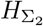. *If H_u_* ⊂ *H*_*l*_, *then* Σ_2_ *is more complex than* Σ_1_.

### Proof

Let 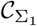 and 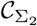 be the complexity classes of Σ_1_ and Σ_2_ respectively. By hypothesis, 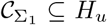 and 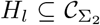 Since, *H*_*u*_ ⊂ *H*_*l*_, we have 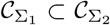 From Lemma 1, it now follows that Σ_2_ is more complex than Σ_1_. □

Indeed, this suggests an economical way to prove complexity results, since the upper bounds and lower bounds could apply to several sets of networks.

In the next section, we apply these notions to explicitly construct a transformation hierarchy and prove some lower bounds for some architectures according to the hierarchy.

## 4 Lower Bounds on the Hierarchy Classes of some Architectures in a certain Transformation Hierarchy

In this section, we construct specific sequences of subsets of F_*m*_ and show that they constitute a transformation hierarchy. Next, we establish some lower bounds on the hierarchy classes of some architectures in this hierarchy. To prove that a certain set in a hierarchy is a lower bound on a hierarchy class, it suffices to show a transformation that is not in the set, yet is in the complexity class in question.

We start off by defining a class of transformations parameterized by a positive integer. For the sake of exposition, we will start by defining a *First-order Transformation* which we will then generalize to a *k*^*th*^-*order Transformation*. We will then show that for all *j* ≥ *I* every *i*^*th*^ order transformation is also a *j*^*th*^ order transformation.

Intuitively, a first-order transformation has the flavor of an SRM_0_ neuron model [30], in that each synapse has a “kernel” function such that effects of inputs spikes according to this kernel are summed over all input spikes across all synapses. The transformation prescribes an output spike if and only if this sum equals a certain “threshold”, which is a positive number.

### Definition 4 (First-order Transformation)

A transformation 𝒯: ℱ_*m*_ →*𝒮* is said to be a *First-order Transformation* if there exists a *τ* ∈ ℝ^+^ and functions *f*_*j*_: ℝ →ℝ, for 1 ≤ *j* ≤ *m*, so that for every *χ* ∈ ℱ_*m*_ and *t* ∈ ℝ, we have Ξ_*t*_𝒯 (*χ*) = ⟨*t*⟩ if and only if we have 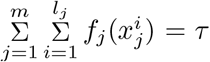 where 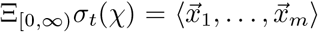 with 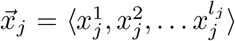 for 1 ≤ *j* ≤ *m*.

Informally, a *k*^*th*^-order transformation is a generalization of a first-order transformation with higher-dimensional kernel functions. Thus, a second-order transformation, for example, has functions that take every pair of spikes and add up their “effects”, in addition to “first-order” effects.

### Definition 5 (*k*^*th*^-order Transformation)

A transformation 𝒯:ℱ_*m*_ →*𝒮* is said to be a *k*^*th*^-*order Transformation* if there exists a *τ* ∈ ℝ^+^ and functions *f*_*j*1_: ℝ → ℝ, *f*_*j*1,*j*2_: ℝ^2^ → ℝ,…, *f*_*j*1 *j*2,.*jk*_: ℝ^*k*^ → ℝ, with 1 ≤ *j*_*p*_ ≤ *m*, where 1 ≤ *p* ≤ *k*, so that for every *χ* ∈ ℱ_*m*_ and *t* ∈ ℝ, we have Ξ 𝒯 (*χ*) = ⟨*t*⟩ if and only if we have 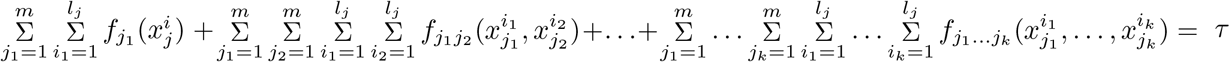 where 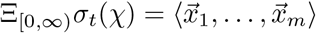 with 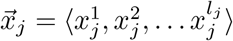, for 1 ≤ *j* ≤ *m*.

For all *j* ≥ *i*, every *i*^*th*^-order transformation is also a *j*^*th*^-order transformation. This can be seen by setting the value of all the functions whose domain has dimensionality greater than *i* to be zero everywhere. Therefore, for all *j* ≥ *i*, the set of all *i*^*th*^-order transformations is a subset of the set of all *j*^*th*^-order transformations. This naturally induces a transformation hierarchy in ℑ_*m*_

### Proposition 1

*For k* ∈ ℤ^+^, *let* 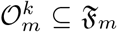 *be the set of all k^th^-order transformations of the form* 𝒯:ℱ_*m*_ →𝒮. *Then,* 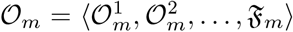 *is a transformation hierarchy in* ℑ_*m*_

For certain acyclic network architectures, we now establish some lower bounds on their hierarchy classes in the above-mentioned transformation hierarchy.

### Theorem 1

*Let* Σ _2_ *be the set the set of all networks with the architecture of the network in Figure 5. Then* 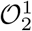 *is a lower bound on the hierarchy class of* Σ _2_ *in* 𝒪_2_.

**Figure 5.**
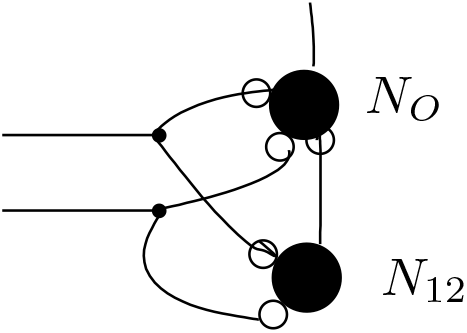
Diagram depicting architecture of networks in Σ_2_.

### Proof

We prove that 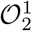 is a lower bound on the hierarchy class of Σ _2_ in 𝒪_2_ by showing a transformation that a network in Σ _2_ can effect, but which lies outside 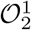.

For the sake of brevity, we describe the salient responses of the neurons from which it is straightforward to construct an SRM_0_ model for them. For the sake of contradiction assume that the transformation effected by the network is a first-order transformation. In Figure 5, neuron *N*_12_ is an inhibitory neuron and the neuron *N*_0_ is an excitatory neuron. Both the input spike trains provide excitatory input to both neurons. We assume, for the sake of contradiction, that the potential of the output neuron *N*_0_ can be written down as a first-order transformations. The argument is made on two input spikes that occurred *t*_1_ and *t*_2_ seconds ago in the first and second input spike train respectively.Consider the values of the functions *f*_1_(*t*_1_) and *f*_2_(*t*_2_), where *f*_1_(·) and *f*_2_(·) are component functions of the putative first-order function in the transformation. The neuron *N*_0_ is set up so that it produces a spike now, if a spike happens either at *t*_1_ alone or *t*_2_ alone. Since the transformation is firstorder, this gives us two equations *f*_1_(*t*_1_) = *τ* and *f*_2_(*t*_2_) = *τ.* When there is a spike both at positions *t*_1_ and *t*_2_, *N*_0_ would reach threshold earlier. However, the occurrence of both these spikes causes *N*_12_ to spike. This, in turn, causes an inhibitory effect on the membrane potential of *N*_0_, which compensates for the extra excitation, so that it spikes exactly once, now. We therefore have the equation *f*_1_(*t*_1_) + *f*_2_(*t*_2_) = *τ.* These three equations (in the two variables *f*_1_(*t*_1_) and *f*_2_(*t*_2_)) form an inconsistent system of linear equations, and therefore *f*_1_(*t*_1_) and *f*_2_(*t*_2_) do not exist, contradicting our hypothesis. Therefore the transformation induced by the current network is not a first-order transformation. Thus, 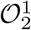 is a lower bound on the hierarchy class of Σ _2_ in 𝒪 _2_.□

Next, we apply a similar strategy to derive a lower bound on the hierarchy class of another network architecture. It will then be clear how one can generalize the present technique.

### Theorem 2

*Let* Σ _3_ *be the set the set of all networks with the architecture of the network in Figure 6. Then* 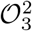 *is a lower bound on the hierarchy class of* Σ _3_ *in* 𝒪 _3_.

**Figure 6.**
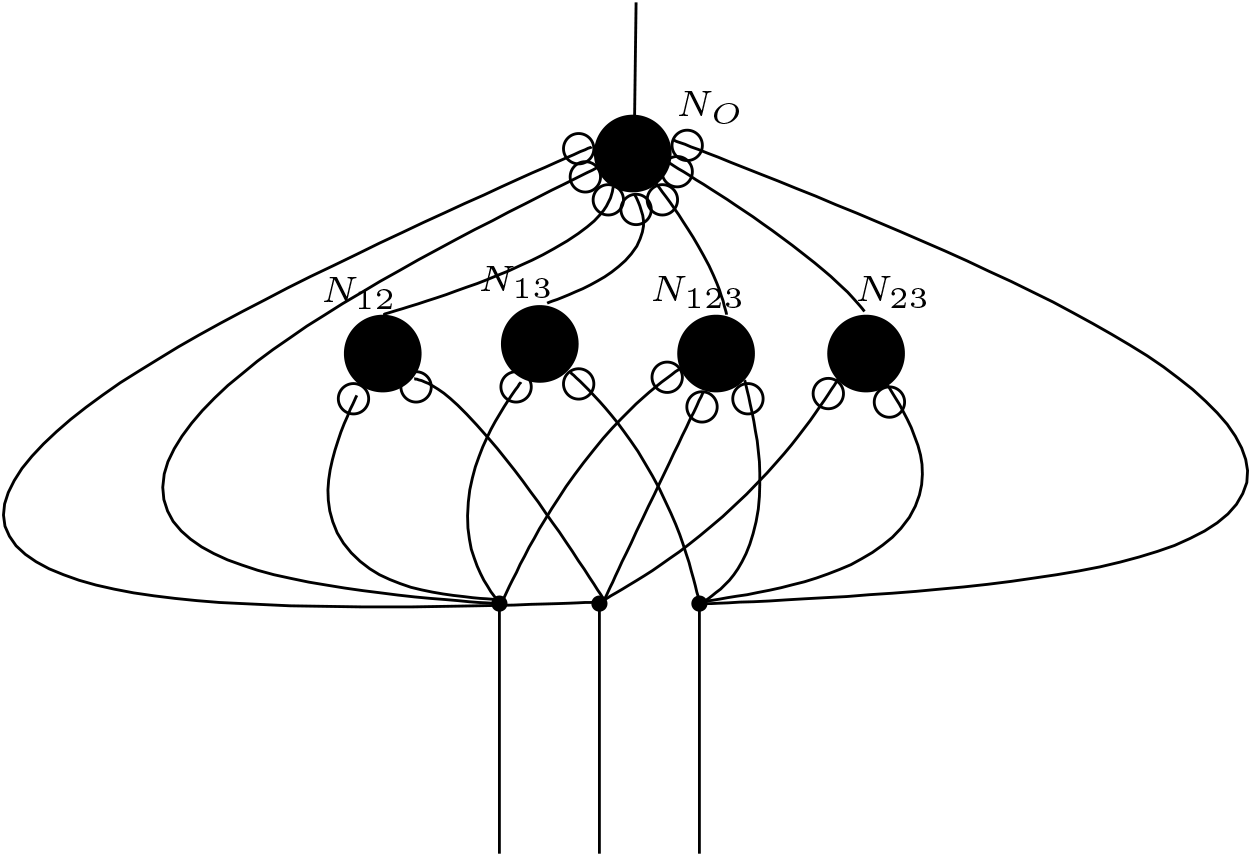
Diagram depicting architecture of networks in Σ_3_.

### Proof

As before, we prove this by showing a transformation that a network in Σ _3_ can effect, but which lies outside 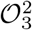. The argument is made on three spike positions *t*_1_, *t*_2_ and *t*_3_ in the past, and the values of the functions *f*_1_(*t*_1_), *f*_2_(*t*_2_), *f*_3_(*t*_3_), *f*_12_(*t*_1_, *t*_2_), *f*_23_(*t*_2_, *t*_3_) and *f*_31_(*t*_3_, *t*_1_), which are component functions of the putative second-order transformation. Again, *N*_0_ is set up so it spikes on each of the three individual spikes occurring alone. This gives us the equations *f*_1_(*t*_1_) = *τ*, *f*_2_(*t*_2_) = *τ* and *f*_3_(*t*_3_) = *τ. N*_12_ works exactly as in the previous example, and so do *N*_23_ and *N*_31_, so as to make the output neuron spike whenever every pair of spikes occur. This gives us the equations *f*_1_(*t*_1_) + *f*_2_(*t*_2_) + *f*_12_(*t*_1_, *t*_2_) = *τ*, *f*_2_(*t*_2_) + *f*_3_(*t*_3_) + *f*_23_(*t*_2_, *t*_3_) = *τ* and *f*_3_(*t*_3_) + *f*_1_(*t*_1_) + *f*_31_(*t*_3_, *t*_1_) = *τ.* Now, when spikes occur simultaneously at all three times, the inhibition provided by *N*_12_, *N*_23_ and *N*_23_ causes the membrane potential of *N*_0_ to always stay below threshold. The neuron *N*_123_ now provides enough excitation to *N*_0_, in order to make it spike now. This gives us the equation*f*_1_(*t*_1_) + *f*_2_(*t*_2_) + *f*_3_(*t*_3_) + *f*_12_(*t*_1_, *t*_2_) + *f*_23_(*t*_2_, *t*_3_) + *f*_31_(*t*_3_, *t*_1_) = *τ.* It is straightforward to verify that this system of 7 equations in 6 variables is inconsistent. Therefore, 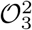 is a lower bound on the hierarchy class of Σ _3_ in 𝒪 _3_.□

It is straightforward to obtain similar results for higher-order transformations with this broad technique.

## 5 Discussion

This work is part of an ongoing research program to build first-principles theory for Connectomics. The idea is to have a theoretical understanding of how structure of neuronal networks constrains spike-timed computations performed by them. This could be useful in generating hypotheses about mechanistic computation in networks when one has connectomic information. Indeed, the field of (microscale) Connectomics is scaling up in its ability to produce increasingly large datasets in diverse organisms and such theory is needed in the context of such data. More fundamentally, the theory will also enable us to understand what structural properties of a network are crucial in enabling it to effect particular classes of spike-timed computations. This has potential application in Comparative Connectomics, an emerging sub-field, where the goal is to compare connectomes either of different individuals of the same species or across individuals of different species in order to determine which structural aspects of connectivity are preserved or altered.

Here, our focus has been on studying the space of spike-train to spike-train transformations. We define notions that allow us to carve up this space into nested sequences of subsets and ask if we can encapsulate the subset of transformations spanned by specific network architectures within sets in this hierarchy. Even if we cannot determine the exact hierarchy classes, establishing bounds on them would be progress. Indeed, we explicitly establish a class of such hierarchies and establish lower bounds corresponding to certain classes of network architectures with respect to said hierarchies.

There are at least two reasons for relating complexity classes to transformation hierarchies. First, complexity classes themselves seem to be hard to characterize succinctly in terms of properties of transformations they contain. Instead, we try to understand complexity classes of network architectures relative to these sets in the hierarchy which are easier to characterize using mathematical properties of transformations. The second reason for this approach is that it provides us another – and a possibly more wholesale – way to prove complexity results via bounds on the corresponding hierarchy classes.

## Competing interests

The author declares that he has no competing interests.

## Author’s contributions

The author performed the research and wrote the paper.

## Acknowledgements

The author wishes to thank Arunava Banerjee for discussions. The work was supported in part by the Simons Foundation and in part by a US National Science Foundation grant (NSF IIS-0902230) to Arunava Banerjee. Part of the work appears in the author’s Ph.D. dissertation at the University of Florida.

## Footnotes

[1] The assumptions manifest as axioms in our abstract model.

[2] Formally, the bounds are with respect to the partial ordering induced by set inclusion.

[3] We assume a single fixed absolute refractory period for all neurons, for convenience, although our results would be no different if different neurons had different absolute refractory periods.

[4] In fact, each feedforward network induces a transformation even in the case when input to it is a stream of spikes from a biologically-relevant spiking regime (not preceded by quiescence). The theory towards this is developed in [1]. It turns out (as exposited in [1] that one can work with ℱ_*m*_ without loss of generality in so far as establishing complexity results is concerned.

[5] The ordering is by set inclusion.

[6] Again, formally, the bounds are with respect to the partial ordering induced by set inclusion.

[7] That is, depending on the hierarchy in question, it is possible that both sets have the same hierarchy class, yet one is more complex than the other.

